# Agreement and uncertainty among climate change impact models: A synthesis of sagebrush steppe vegetation projections

**DOI:** 10.1101/2020.06.16.154989

**Authors:** Scott N. Zimmer, Guenchik J. Grosklos, Patrick Belmont, Peter B. Adler

## Abstract

Ecologists have built numerous models to project how climate change will impact rangeland vegetation, but these projections are difficult to validate, making their utility for land management planning unclear. In the absence of direct validation, researchers can ask whether projections from different models are consistent. High consistency across models based on different assumptions and emission scenarios would increase confidence in using projections for planning. Here, we analyzed 42 models of climate change impacts on sagebrush (*Artemisia tridentata* Nutt.), cheatgrass (*Bromus tectorum* L.), pinyon-juniper (*Pinus* L. *spp*. and *Juniperus* L. *spp*.), and forage production on Bureau of Land Management (BLM) lands in the United States Intermountain West. These models consistently projected the potential for pinyon-juniper declines and forage production increases. In contrast, cheatgrass models mainly projected no climate change impacts, while sagebrush models projected no change in most areas and declines in southern extremes. In most instances, vegetation projections from high and low emissions scenarios differed only slightly.

The projected vegetation impacts have important management implications for agencies such as the BLM. Pinyon-juniper declines would reduce the need to control pinyon-juniper encroachment, and increases in forage production could benefit livestock and wildlife populations in some regions. Sagebrush conservation and restoration projects may be challenged in areas projected to experience sagebrush declines. However, projected vegetation impacts may also interact with increasing future wildfire risk in ways single-response models do not anticipate. In particular, forage production increases could increase management challenges related to fire.

## Introduction

### Background

Ecological impacts of climate change, including shifts in species distribution, abundance and phenology, have been widely documented across ecosystems and are predicted to intensify (Parmesan 2006; Parmesan and Yohe 2003; Urban 2015). However, forecasting specific impacts of climate change on given ecosystems remains extremely difficult. Predictive modeling techniques are commonly used to project such climate change impacts, but these projections are rarely validated against direct observations. Whereas the accuracy and uncertainty of short-term forecasts can be assessed, decade-to-century-scale projections most useful in climate change planning are extremely difficult to validate robustly (Araújo and Guisan 2006). As a result, the accuracy of long-term projections concerning ecological impacts of climate change is largely unknown (Knutti 2008), raising questions about their value for land management and planning (Littell et al. 2011; Mouquet et al. 2015; Robinson et al. 2008; Yates et al. 2018).

In the absence of direct model validation, multiple models can be compared to evaluate consistency among their projections. If models reach similar conclusions despite differences in methods, input data, or assumptions, projections become more credible, uncertainty is reduced, and decision-makers may be more confident considering them in planning. Such model intercomparison is a common approach among climate models. The Coupled Model Intercomparison Project (CMIP) (Eyring et al. 2016) and Ocean Model Intercomparison Project (OMIP) (Griffies et al. 2016) are notable model efforts synthesizing general circulation models (GCMs) and earth systems models to better project future climate changes. However, similar efforts focused on modeling the ecological impacts of climate change are rare (Krysanova and Hattermann 2017). Renwick et al. (2018) modeled climate change impacts on sagebrush through four approaches to evaluate consistency across models. Others have used multiple methods to model climate change impacts on stream fish in France (Buisson et al. 2010) and vegetation in California (Crimmins et al. 2013). However, such comparisons remain the exception more than the rule.

Meta-analysis, in which the results of independent studies are compared (Arnqvist and Wooster 1995), offers another form of model comparison especially well-suited for synthesizing results from multiple research groups and eliminating the potential bias of a single group conducting analyses. However, we know of only one climate change impact model meta-analysis focused on natural resources, a study which assessed overall ecosystem-level vulnerability of marine environments to climate change (Queirós et al. 2016).

Here, we synthesized projections from many models of climate change impacts on four keystone vegetation components of the United States Intermountain West–sagebrush (*Artemisia tridentata* Nutt.), cheatgrass (*Bromus tectorum* L.), pinyon-juniper (*Pinus* L. *spp*. and *Juniperus* L. *spp*.) and forage production. We compared model results using a spatially explicit approach, revealing instances of high and low agreement in projected impacts throughout the Intermountain West. We intend for this unique synthesis of climate change impact models to inform land management by addressing individual vegetation components at scales relevant to land managers, incorporating results from multiple researchers, and explicitly addressing uncertainty in projected impacts.

### BLM Land and Vegetation

We conducted this analysis specifically to address climate change impacts on Bureau of Land Management (BLM) lands within the United States Intermountain West. The BLM manages approximately 142 million acres in this region, and has a “multiple use mandate” to administer land for livestock grazing, hunting, recreation, energy extraction, and other land uses (Bureau of Land Management 2016). Vegetation is the foundation for many of these uses, providing forage for domestic livestock and wildlife, habitat for wildlife, and other ecosystem services including protecting soils from erosion, sequestering carbon, and cycling nutrients (Havstad et al. 2007; Yapp et al. 2010).

Major land management issues exist in this area, including the spread of invasive species such as cheatgrass (Bradley 2009; Knapp 1996), encroachment of conifers such as pinyon-juniper into low elevation habitats (Weisberg et al. 2007), and declines in sagebrush (Bradley 2010) which affect fire, grazing management, and wildlife habitat (Beschta et al. 2013). There is also difficulty balancing grazing and other land uses while maintaining habitat quality (Camp et al. 2014). The sagebrush-obligate sage-grouse (*Centrocercus urophasianus*) has received much recent attention, as suitable habitat declines increasingly threaten its populations (Creutzburg et al. 2015). Wildfire is a significant process in sagebrush steppe ecosystems, which can limit establishment of woody perennials (Davies et al. 2012) and create positive feedbacks with cheatgrass (Balch et al. 2013). Understanding how climate change will increasingly interact with these issues is crucial for land management planning.

### Research Objectives

Though various peer-reviewed studies have modeled climate change impacts to vegetation components of the Intermountain West, we know of no efforts to compare their results to synthesize findings and assess agreement between studies. We aim to do such a synthesis to address the following objectives:

1. Review the projected impacts of climate change on sagebrush, pinyon-juniper, cheatgrass, and forage production in the U.S. Intermountain West, according to current vegetation models.
2. Assess whether projected impacts are consistent across models, and whether differences in emissions scenarios influence projected impacts.
3. Interpret the management implications of projected vegetation impacts.

## Methods

### Study Area

Our research was focused on BLM land in the U.S. Intermountain West, defined as the region between the eastern slopes of the Rocky Mountains and the eastern slopes of the Sierra Nevada and Cascade Mountains, stretching from the Mexican to Canadian border. BLM land in this region encompasses approximately 142 million acres across Washington, Oregon, California, Idaho, Nevada, Utah, Arizona, Montana, Wyoming, Colorado and New Mexico, comprising more than one-quarter of the total area of the Intermountain West (Fig. 1). The Intermountain West includes 18 EPA level III ecoregions (US EPA 2015), but BLM land falls predominantly in only four—the Northern Basin and Range (NBR), Central Basin and Range (CBR), Wyoming Basin (WB), and Colorado Plateau (CP). Since these regions are most significant to the management of Intermountain West BLM land, we focused only on those regions.

**Figure 1:**
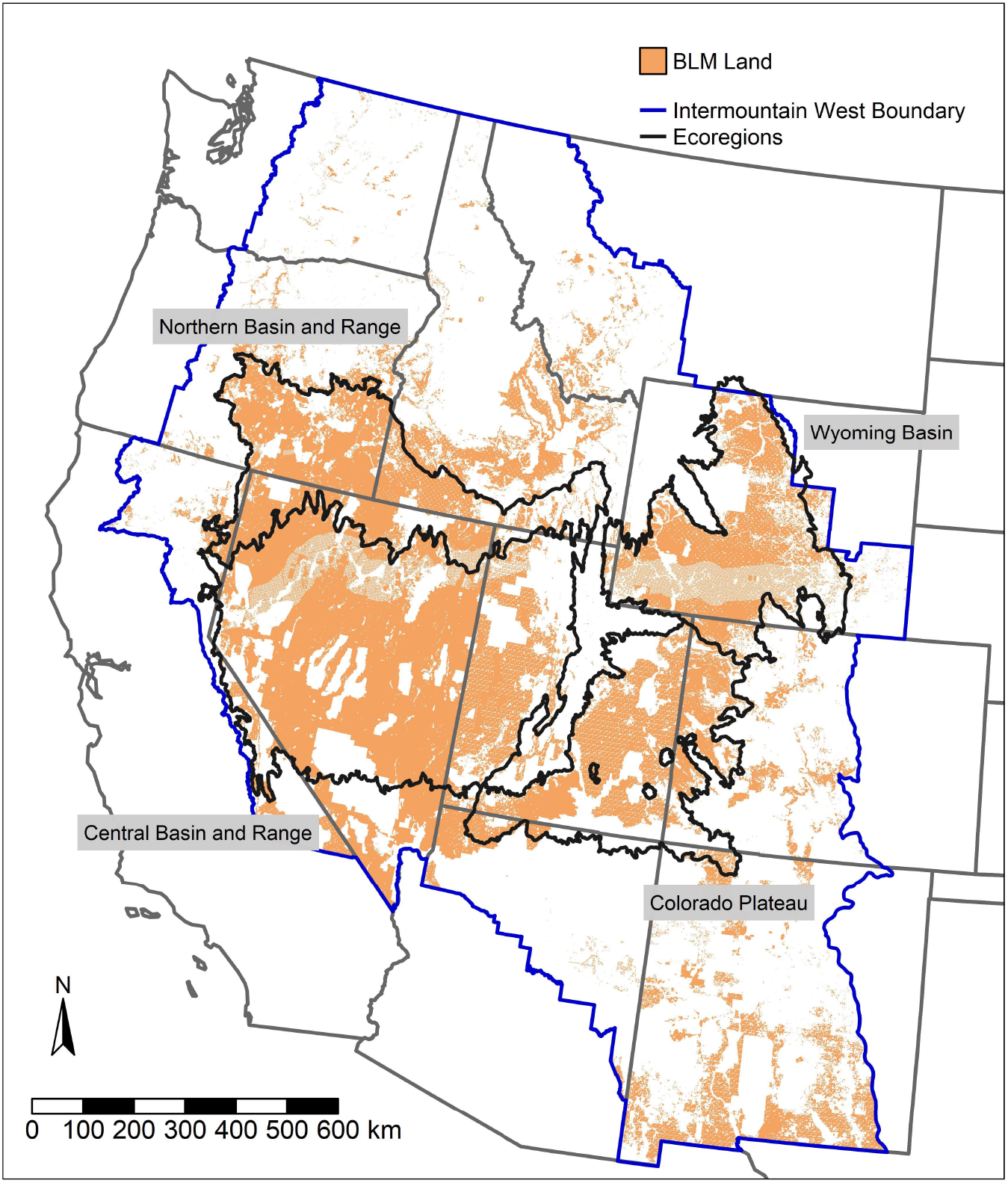
Map of BLM lands within the United States Intermountain West. The four ecoregions containing the majority of BLM land in the Intermountain West are outlined and labeled.

### Obtaining Vegetation Model Projections

We first conducted a literature review by searching Google Scholar for peer-reviewed studies modeling climate change impacts on sagebrush, cheatgrass, pinyon-juniper and forage production in the Intermountain West. We searched each vegetation type in conjunction with the phrase “climate change”, as well as “projections”, “impacts”, “models”, and “forecasts” to find such studies. We selected 14 studies published since 2008 that provided spatially-explicit results of projected impacts to these vegetation types due to climate change. The time constraint was selected to eliminate studies using outdated GCMs or methods. For each model, we noted important metadata such as the model type, emissions scenarios considered, response variables, and the latest time to which results were modeled, since these characteristics may affect comparability of model projections. Spatial extents of the studies differed, but all studies addressed lands within the Intermountain West.

In all, we identified three studies addressing sagebrush (Renwick et al. 2018; Schlaepfer et al. 2012; Still and Richardson 2015), three addressing cheatgrass (Boyte et al. 2016; Bradley 2009; Brummer et al. 2016), five addressing pinyon-juniper (Cole et al. 2008; Jiang et al. 2013; McDowell et al. 2016; Notaro et al. 2012; Rehfeldt et al. 2012), and three addressing forage production (Hufkens et al. 2016; Notaro et al. 2012; Reeves et al. 2017). Many studies made multiple projections, employing either different emissions scenarios or model types. From the 14 studies, we obtained 42 distinct projections of change for these vegetation components. Metadata for these studies are shown Table 1.

**Table 1:**
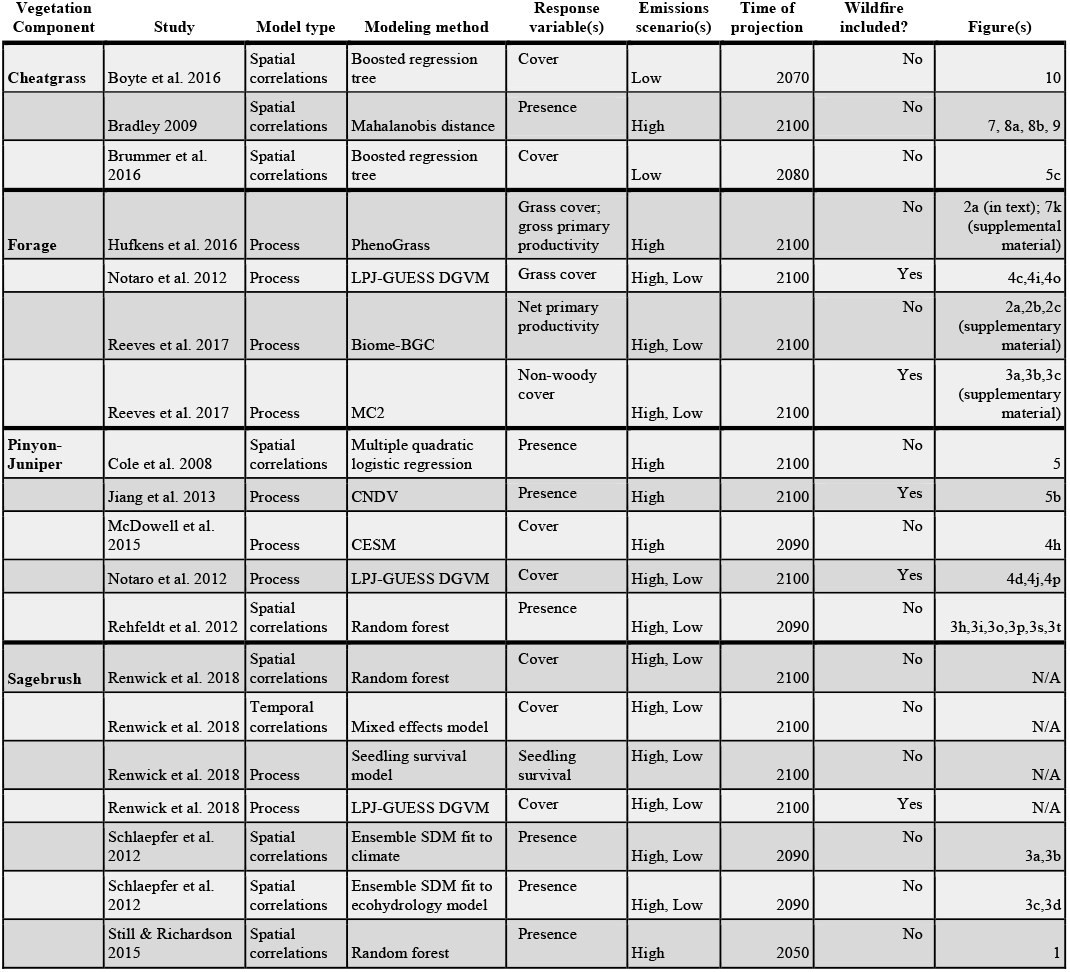
Studies from which model results were analyzed, showing metadata for model comparison. A1b, A2 and RCP 8.5 emissions scenarios were considered high emissions. B1, B2, and RCP 4.5 were considered low emissions. Results from Renwick et al. 2018 are supplemental results obtained from authors.

The models we found can be broadly categorized as either correlations-based, including spatial and temporal correlations, or process-based. Spatial correlation models correlate current species distributions or abundances to current climatic and environmental conditions, and predict future distribution based on projected future climate (Elith and Leathwick 2009). Temporal correlations models relate the effects of current interannual climatic variation to ecological responses, and project these relationships using future climate (Kleinhesselink and Adler 2018). Process-based models consider the mechanistic processes driving system dynamics in order to project impacts of changing climate (Johnsen et al. 2001; Larocque et al. 2015). More simply, process models explicitly define mechanistic model parameters, whereas these mechanisms are implicit in correlations models (Dormann et al. 2012).

The process-based models included in our study varied widely, and utilized numerous distinct model programs, such as PhenoGrass (Hufkens et al. 2016), BIOME-BGC (Reeves et al. 2017), MC2 (Reeves et al. 2017), CNDV (Jiang et al. 2013), CESM (McDowell et al. 2016), and LPJ-GUESS (Notaro et al. 2012; Renwick et al. 2018). The spatial and temporal correlation models we used projected climate-related vegetation changes using random forest (Rehfeldt et al. 2012; Renwick et al. 2018; Still and Richardson 2015), boosted regression trees (Boyte et al. 2016; Brummer et al. 2016), Mahalanobis distance (Bradley 2009), or ensembles (Schlaepfer et al. 2012).

Since forage represents vegetation available for grazing, rather than a particular species, forage was modeled differently depending on the study. Forage production can be represented as grassland cover (Hufkens et al. 2016; Notaro et al. 2012) or relative abundance of non-woody vegetation (Reeves et al. 2017), but we also included two studies which modeled changes to primary productivity (Hufkens et al. 2016; Reeves et al. 2017), which can be interpreted as a change in forage quantity. Increases in primary productivity do not necessarily translate directly to forage increases, but unless increasing primary productivity comes with a loss of palatable vegetation, it indicates an increase in biomass available for grazers and therefore a likely increase in forage quantity.

### Analyzing Vegetation Model Projections

We analyzed the results of models in a spatially-explicit framework by downloading the highest resolution image available for each figure, importing those into ArcMap, and georeferencing them. Georeferencing applies a geographic projection to an image and aligns it with the area to which it corresponds by anchoring it at easily identifiable features like state borders. This process converted the original image into a raster whose resolution was determined by the resolution of the original image. We then conducted an unsupervised classification in R (Hijmans 2018; R Core Team 2018) to identify pixel groups in each raster, and manually assigned these pixel groups values of −1. 0, or 1, corresponding to a projected decrease in the vegetation component, no change, or an increase, respectively.

For example, if a given image visualized a decrease in sagebrush abundance as red pixels, the classification collected all red pixels in the image into a group, which we assigned a value of −1. This allowed us to only analyze the direction of vegetation impacts, not the magnitude of impacts. This limitation was acceptable because many models we included analyzed changes in presence/absence of vegetation (Table 1), for which magnitude of change cannot be determined. Pixels not corresponding to vegetation impacts, such as background, text and legends were removed by assigning them a value of NA. Every raster was masked by a polygon layer of BLM land in the Intermountain West (U.S. Geological Survey 2017) to eliminate unnecessary information.

For each vegetation component, we then resampled all rasters by nearest neighbor to the finest pixel resolution and stacked all rasters. Lastly, we counted the number of models indicating an increase, decrease, or no change at each pixel for each vegetation type. We only considered pixels where at least three models made projections. We then visualized the counts of models projecting decreases, increases, or no change as RGB images. In these images, the number of models indicating a decrease in a given vegetation type determined the red intensity, number of models indicating an increase determined green intensity, and number of models indicating no change determined blue intensity. This visualization gave a broad sense of model projections on a pixel-by-pixel basis, and revealed where models consistently projected a particular outcome and instances where projections were split between multiple outcomes. This visualization was complemented by a legend showing the mean RGB value within each ecoregion (Smith 2017). An example of how RGB values were plotted onto the legend is shown in Fig. S1. We made these visualizations showing all results for a given vegetation type, as well as results split by emissions scenario.

We did not assess the statistical significance of these outcomes for a number of reasons. First, the RGB maps described above communicate all the information in the original studies about degree of agreement in the direction of change at the resolution of individual pixels. It is not clear what additional information a *p*-value would add. Second, while it is technically possible to test statistical-significance at the level of individual pixels (e.g., Chi-square or multinomial), the results have low power due to small sample sizes, as small as 3 model projections at a given pixel after separating results from high and low emissions scenarios. In addition, pixel-level tests would ignore spatial autocorrelation. We believe that the descriptive, visual results we produced are easier to interpret than a potentially flawed statistical analysis.

We considered only the most long-term future projections in each study. Most studies projected through 2090 or 2100, though one projected changes only to 2050. Shorter term projections may indicate lesser climate change impacts than longer term projections, but the direction of change is unlikely to vary between these time frames. Since we considered only direction of change, not magnitude, the impact of varying projection times should be minimal.

## Results

We produced RGB visualizations revealing the overall projected impacts for each vegetation component on a pixel-by-pixel basis (Figs. 2–5). These results were split to show results among models employing low emissions scenarios, high emissions scenarios, and across all emissions scenarios.

**Figure 2:**
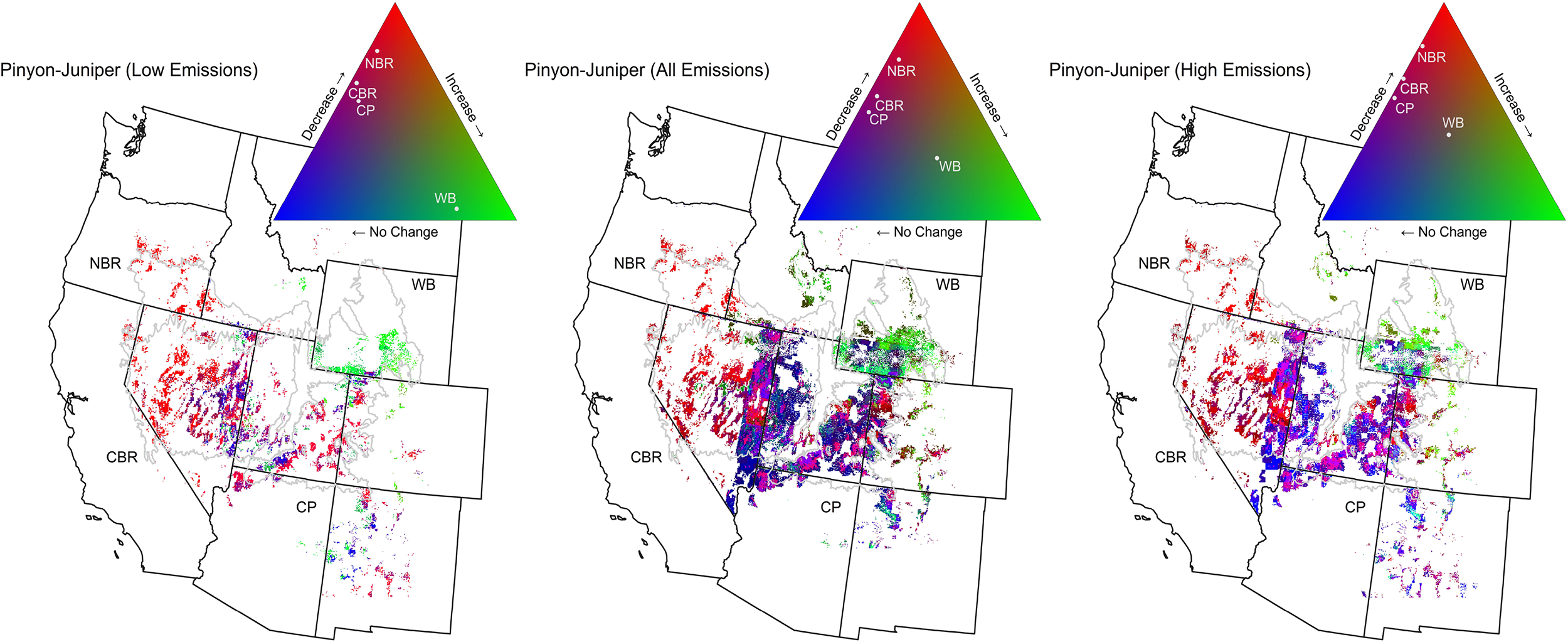
RGB visualizations for pinyon-juniper, among models employing low emissions scenarios, all emissions scenarios, and high emissions scenarios. Red intensity is determined by the number of models indicating a decrease in vegetation, green intensity is determined by number of models indicating an increase, and blue intensity is determined by number of models indicating no change. Muted colors indicate greater inconsistencies in projected direction of change. Mean pixel values of each ecoregion are shown (white points with text labels) within the legends for each emission scenario.

**Figure 3:**
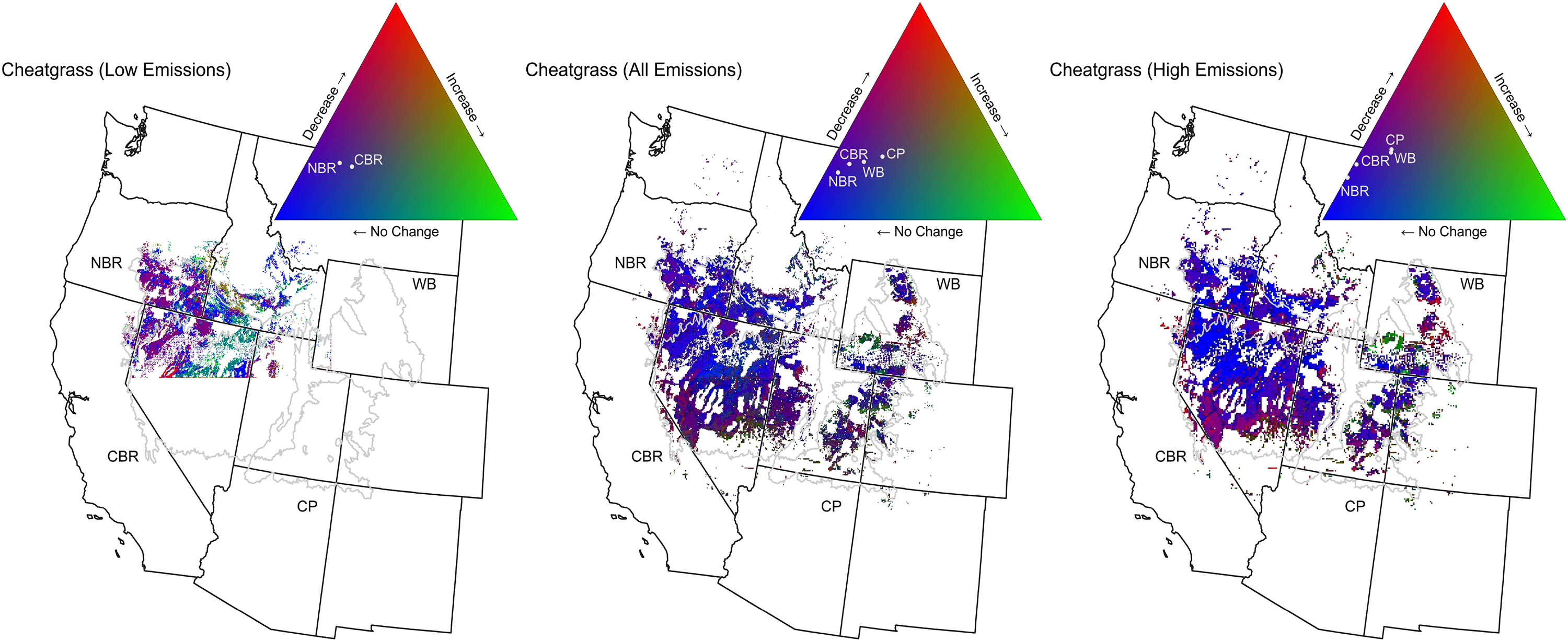
RGB visualizations for cheatgrass, among models employing low emissions scenarios, all emissions scenarios, and high emissions scenarios. Only two included models made cheatgrass projections under low emissions scenarios, so we included pixels addressed by both of those models. Results are not available for CP and WB because both models did not address these regions.

**Figure 4:**
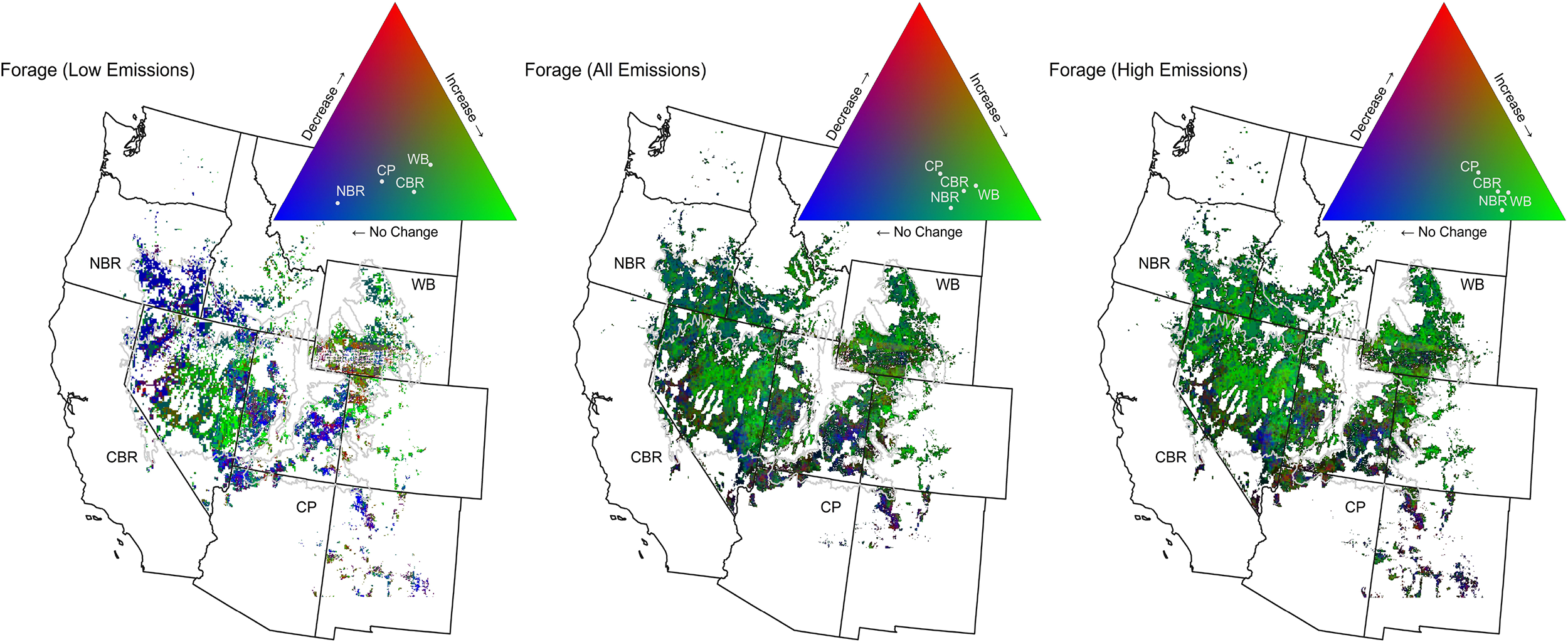
RGB visualizations for forage production, among models employing low emissions scenarios, all emissions scenarios, and high emissions scenarios.

**Figure 5:**
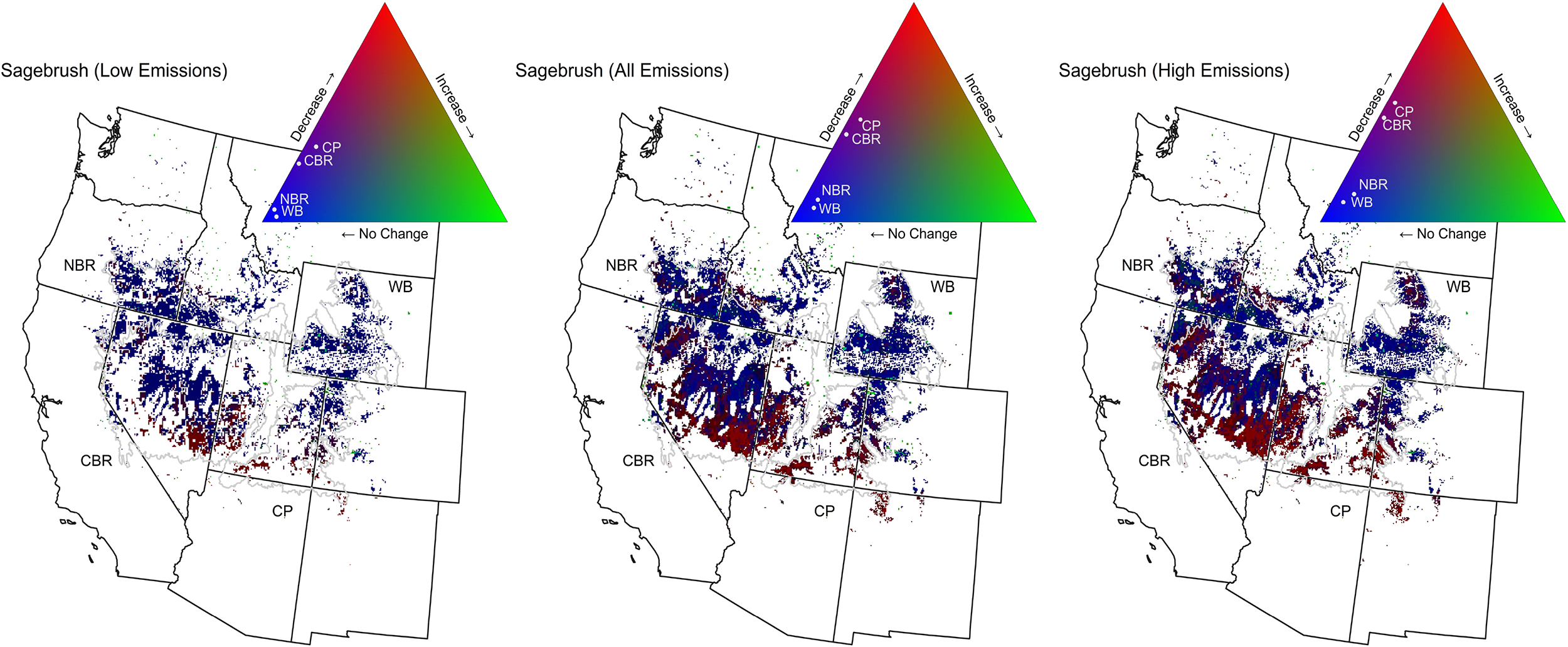
RGB visualizations for sagebrush, among models employing low emissions scenarios, all emissions scenarios, and high emissions scenarios.

Pinyon-juniper projections showed decreases in all ecoregions except for WB (Fig. 2). These decreases were strongest in NBR, and were consistent across emissions scenarios. Results in CBR and CP were also consistent across emissions scenarios, and mainly showed decreases along with some indications of no change. Emissions scenario had the greatest influence on pinyon-juniper projections in WB, with low emissions scenarios showing increases, and high emissions results showing highly uncertain results mixed between increases, decreases, and no change. While results were highly uncertain in this region as a whole, it should be noted that smaller areas within it more consistently indicate the potential for increases.

Cheatgrass results tended toward no change in all ecoregions, and were consistent across emissions scenarios (Fig. 3). In NBR and CBR, both low and high emissions results mainly showed indications of no change, though low emissions results were slightly less certain than high emissions results in these regions. Results in CP and WB were more uncertain or showed varying directions of change, but still tended toward no change. There were insufficient models employing low emissions scenarios within CP and WB, so low emissions results were not available in these regions.

Forage production results varied by emissions scenario, with high emissions results showing consistent indications of increase in all ecoregions, while low emissions results were more uncertain in general (Fig. 4). In WB and CBR, low emissions results also tended toward increases, but with somewhat more uncertainty than high emissions results. On the other hand, low emissions results conflicted with high emissions results in NBR and CP. Whereas high emissions results showed increases in these regions, low emissions results indicated no change in NBR, and in CP were split between no change and increases.

Sagebrush results were consistent across emissions scenarios, showing either no change or decreases (Fig. 5). In WB and NBR, results consistently indicated no change in sagebrush for both high and low emissions scenarios. Results were more uncertain in CBR and CP, split between no change and decreases. Among low emissions scenarios, results tended more toward no change in these ecoregions, whereas high emissions results tended more towards decreases. Latitude seemed to strongly affect these results, as southern areas of these regions indicated decreases regardless of emissions scenario and northern areas consistently showed no change.

Our sagebrush analysis included non-gridded, point projections from one study with eight models (Renwick 2018). To include these points in our analysis, we converted them to individual raster cells, but these covered little area and thus had little weight. When we considered only the cells included in Renwick et al. results differed slightly (Fig. S2), suggesting sagebrush may increase in many places where Fig. 5 indicated no change. However, the Renwick et al. subset was consistent with the full analysis in indicating sagebrush declines in its southern range.

## Discussion

### Projected Vegetation Changes

For pinyon-juniper, we mainly found indications of decreases with some possibility of no change within the Northern Basin and Range, Central Basin and Range, and Colorado Plateau (Fig. 2). Results did not strongly vary by emissions scenario. Reviews have argued that conifers such as pinyon pine and juniper may face widespread mortality in the future (Allen et al. 2015; van Mantgem et al. 2009), and our results largely support this conclusion in all ecoregions except the Wyoming Basin, where projections indicate increases among low emissions scenarios and are uncertain among high emissions scenarios. The Wyoming Basin is a distinct spur of sagebrush steppe with a cool/moist soil moisture and temperature regime (Chambers et al. 2014), so increasing temperatures and precipitation changes may have unique effects which benefit pinyon-juniper more here.

Projected impacts of climate change on cheatgrass were weak in the Northern and Central Basin and Range, and uncertain in the Colorado Plateau and Wyoming Basin, indicating the possibility of multiple outcomes (Fig. 3). Some experimental studies have suggested cheatgrass can benefit from climate change effects at local scales (Compagnoni and Adler 2014; Zelikova et al. 2013; Ziska et al. 2005), somewhat at odds with the models we reviewed. Cheatgrass is strongly associated with local disturbances (Bradford and Lauenroth 2006) that may also be related to climate change, such as wildfire (Abatzoglou et al. 2017), so the aggregate direct and indirect impacts of climate change on cheatgrass may be difficult to model at regional scales.

Our analysis suggests forage production increases are generally likely in the ecoregions we analyzed (Fig. 4). Indications of forage increases were strongest in the Wyoming Basin and Central Basin and Range. In the Northern Basin and Range and Colorado Plateau, high emissions scenarios showed increases, but low emissions scenario results were more uncertain in the Colorado Plateau and indicated no change in the Northern Basin and Range. Our overarching finding of increasing forage production is consistent with much of the literature (Izaurralde et al. 2011; Polley et al. 2013), which anticipates forage production increases largely as a result of increasing CO_2_ concentrations (Friend, 2014). The more mixed results we found among low emissions scenarios is therefore not surprising.

Sagebrush projections showed little change in the Northern Basin and Range and Wyoming Basin, while projections in the Central Basin and Range and Colorado Plateau were split between decreases and no change (Fig. 5). The decreases were consistently concentrated in the southern portion of these regions, but also in the low elevation Snake River Plain north of the Northern Basin and Range. Studies typically anticipate sagebrush increases in its northern range and high elevations and decreases in its southern range and low elevations (Bradley 2010; Kleinhesselink and Adler 2018). Our results support the projected declines in the south, but suggest little change is likely in northern regions.

### Emissions Scenario Uncertainty

Uncertainty in emissions scenarios, and therefore uncertainty in the degree of climatic changes that will occur in the future, can complicate climate change adaptation planning (Dessai and Hulme 2007). However, we generally found a high degree of agreement in climate change impacts projected by models employing different emissions scenarios. While we were not able to quantitatively assess the magnitude of projected vegetation impacts, which may be more sensitive to emissions scenario, we found that in most cases the direction of projected impacts was not strongly influenced by emissions scenarios in our study area. Therefore, management actions can likely be planned despite uncertainty in future emissions trajectories, as we expect similar shifts for these vegetation types regardless of emissions scenario.

However, we found conflicting results depending on emissions scenarios in a handful of cases. High emissions scenarios indicated forage production increases in all regions, but low emissions scenarios indicated no impacts in Northern Basin and Range and uncertain results in Colorado Plateau. Pinyon-juniper projections in Wyoming Basin were also influenced by emissions scenario, with low emissions scenarios indicating increases, and high emissions scenarios showing uncertain results. In these instances, overall projections should be seen as more uncertain, given the potential for differing outcomes depending on emissions scenario.

### Model Choice Uncertainty

We considered two sources of model choice uncertainty. First, models may differ in approach, such as process-based versus spatial or temporal correlations models. Even among studies using a similar approach, such as spatial correlations, differences in statistical methods can lead to variation in predictions. For example, a comprehensive evaluation of 33 species distribution models found statistical methods strongly influenced predictive performance (Norberg et al. 2019), and others have found fitting method can affect distribution model results more than emissions scenario or GCM (Buisson et al. 2010; Diniz-Filho et al. 2009). Therefore, a consensus in projected impacts from models using contrasting approaches and input data would increase confidence in projections. On the other hand, a consensus in projections from models based on similar approaches is less meaningful, as any single approach may have biases or gaps toward a given result.

Across the ecological responses we studied, there was variation in diversity of approaches and methods. For example, the cheatgrass models we found had no diversity of approach, since all were spatial correlations models, but had some diversity in statistical methods, with two studies modeling cover through boosted regression trees while one modeled presence/absence with Mahalanobis distance (Table 1). This limited ensemble may therefore underestimate model choice uncertainty. Similarly, we found only process-based models of forage production. However, each of these studies utilized a different simulation model (Biome-BGC, MC2, etc.) and response variables also differed between models (grass cover, non-woody vegetation cover, gross primary productivity, and net primary productivity), making this suite of models more diverse. Pinyon-juniper and sagebrush featured the most varied ensembles— process and correlations models were available for each of these vegetation types, and modeling methods and response variables also differed between models. This diversity increases our confidence in the projections. Despite the diversity of these ensembles, pinyon-juniper and sagebrush projections were no more inconsistent than those for cheatgrass or forage production.

### Limitations

There is inherent uncertainty in projecting future climate with GCMs (Deser et al. 2007), and this uncertainty carries over into any models projecting ecological impacts of climate change. These models are complex, but cannot consider all possible drivers of ecological change. For example, increases in extreme weather events due to climate change are likely (IPCC 2007), but may not be explicitly considered by many ecological impact models. Models account for interactions such as species competition or disturbance from wildfire or grazing in various ways, another limitation that can affect their generalizability.

Wildfire in particular may be important to consider in relation to our results. Increases in wildfire size, frequency and intensity in response to climate change are expected in the Intermountain West (Abatzoglou and Williams 2016; Barbero et al. 2015; Murphy et al. 2018; Prudencio et al. 2018) due to warmer, drier conditions and excessive fuel loads (Liu and Wimberly 2016; Murphy et al. 2018). The projected forage production increases we found could intensify these wildfire dynamics as well. In the models we analyzed, forage represents either overall plant productivity or grass cover—increases in these measures indicate more total fuels or more fine fuels, respectively, either of which could promote wildfires.

Wildfire increases could greatly impact the vegetation we considered. Pinyon-juniper and sagebrush are extremely susceptible to fire (Allen et al. 2015; McDowell et al. 2016; Reeves et al. 2018), meaning increased wildfire could encourage greater declines in pinyon-juniper and sagebrush than projected by the models we analyzed. Cheatgrass promotes and benefits from wildfire through the “cheatgrass-fire cycle” (Balch et al. 2013; Bradley et al. 2018), so increased wildfire could benefit cheatgrass beyond what was projected in the models we reviewed (Larson et al. 2018). Only four models we analyzed specifically accounted for wildfire (Jiang et al. 2013; Notaro et al. 2012; Reeves et al. 2017; Renwick et al. 2018), so wildfire interactions are not well accounted for in our results.

## Implications

The potential for pinyon-juniper declines in the Colorado Plateau, Central Basin and Range, and Northern Basin and Range indicate the BLM may be able to reduce pinyon-juniper management actions such as chaining and prescribed burns in some areas. Such management actions have been widespread for decades to control pinyon-juniper encroachment (Redmond et al. 2013). We mainly found indications of no projected climate change impacts to cheatgrass, indicating current cheatgrass management objectives should continue into the future. However, cheatgrass projections are strongly correlated with precipitation amount and seasonality (Bradley 2009), which remain difficult to project into the future. Therefore, cheatgrass projections could change as future precipitation predictions improve. Since other invasive grasses such as medusahead (*Taeniatherum caput-medusae* L.) and red brome (*Bromus rubens* L.) could fill the cheatgrass niche if it were to decline (Snyder et al. 2019), invasive grass management needs are likely to continue, though.

Projected forage increases across our study area could have major management implications, positively impacting livestock grazing and wildlife. However, overall management implications may be more complex, as climate change could also increase heat stress on livestock and wildlife and make forage production more variable (Reeves et al. 2017). Indications of no impacts on sagebrush in many areas indicate current sagebrush management can continue in the future through much of the Intermountain West. We did find the potential for sagebrush declines in southern parts of the Central Basin and Range and Colorado Plateau, though, implying that conservation and restoration investments in these areas may be risky. Wildfire is likely to become more intense and frequent due to climate change, which will also have important direct and indirect management implications interacting with these vegetation impacts.

## Conclusion

We conducted a spatially-explicit synthesis of models projecting climate change impacts to vegetation on BLM lands throughout the Intermountain West. We found many instances where climate change is projected to impact vegetation, and these impacts are generally consistent across high and low emissions scenarios. These projected impacts have clear implications for future management of BLM lands, and the general agreement between high and low emissions results indicates that planning can proceed despite uncertainty in future emissions reductions.

Projections of the ecological impacts of climate change are inherently uncertain, given the complexity of ecological systems and the uncertainty of future climate projections and emissions scenarios, all of which are addressed by researchers through different approaches. Thus, we believe syntheses like ours are useful tools to determine where projections are consistent despite such uncertainties, and may reliably inform policy and management. Analyses like ours are lacking, and would be more feasible if model results could be easily accessed in central databases. For example, Quierós et al. (2016) accessed results from such a database to conduct a meta-analysis of marine climate change impacts. Other efforts exist to collect models of impacts on water, agriculture, and marine environment (Tittensor et al. 2018; Warszawski et al. 2014), but we are unaware of a database focused on vegetation impacts at more local scales. Such a database of spatially-explicit results could be an invaluable resource for other researchers to robustly compare model outputs.

## Acknowledgements

We would like to thank all students and faculty within the Climate Adaptation Science program, for help encouraging our research and refining our ideas. Courtney Flint provided essential support throughout the project, as did Elaine Brice, Kirsten Goldstein, Brett Miller, and Hongchao Zhang. Thank you to Nancy Huntly for continued guidance and support of the Climate Adaptation Science program, and to Hadia Akbar, Rachel Hager, Tara Saley, and Emily Wilkins for feedback on our project.

## Data Availability

The underlying data and code used in this analysis are available online (Zimmer et al. 2019).

## References

Abatzoglou, J.T., Kolden, C.A., Williams, A.P., Lutz, J.A., Smith, A.M.S., 2017. Climatic influences on interannual variability in regional burn severity across western US forests. International Journal of Wildland Fire 26, 269. https://doi.org/10.1071/WF16165

Abatzoglou, J.T., Williams, A.P., 2016. Impact of anthropogenic climate change on wildfire across western US forests. Proceedings of the National Academy of Sciences 113, 11770–11775. https://doi.org/10.1073/pnas.1607171113

Allen, C.D., Breshears, D.D., McDowell, N.G., 2015. On underestimation of global vulnerability to tree mortality and forest die-off from hotter drought in the Anthropocene. Ecosphere 6, art129. https://doi.org/10.1890/ES15-00203.1

Araújo, M.B., Guisan, A., 2006. Five (or so) challenges for species distribution modelling. Journal of Biogeography 33. 1677–1688. https://doi.org/10.1111/j.1365-2699.2006.01584.x

Arnqvist, G., Wooster, D., 1995. Meta-analysis: synthesizing research findings in ecology and evolution. Trends in Ecology & Evolution 10, 236–240. https://doi.org/10.1016/S0169-5347(00)89073-4

Balch, J.K., Bradley, B.A., D’Antonio, C.M., Gómez Dans, J., 2013. Introduced annual grass increases regional fire activity across the arid western USA (1980–2009). Global Change Biology 19, 173–183. https://doi.org/10.1111/gcb.12046

Barbero, R., Abatzoglou, J.T., Larkin, N.K., Kolden, C.A., Stocks, B., 2015. Climate change presents increased potential for very large fires in the contiguous United States. International Journal of Wildland Fire 24, 892. https://doi.org/10.1071/WF15083

Beschta, R.L., Donahue, D.L., DellaSala, D.A., Rhodes, J.J., Karr, J.R., O’Brien, M.H., Fleischner, T.L., Deacon Williams, C., 2013. Adapting to Climate Change on Western Public Lands: Addressing the Ecological Effects of Domestic, Wild, and Feral Ungulates. Environmental Management 51, 474–491. https://doi.org/10.1007/s00267-012-9964-9

Boyte, S.P., Wylie, B.K., Major, D.J., 2016. Cheatgrass Percent Cover Change: Comparing Recent Estimates to Climate Change-Driven Predictions in the Northern Great Basin. Rangeland Ecology & Management 69, 265279. https://doi.org/10.1016/j.rama.2016.03.002

Bradford, J.B., Lauenroth, W.K., 2006. Controls over invasion of *Bromus tectorum□*: The importance of climate, soil, disturbance and seed availability. Journal of Vegetation Science 17, 693–704. https://doi.org/10.1111/j.1654-1103.2006.tb02493.x

Bradley, B.A., 2010. Assessing ecosystem threats from global and regional change: hierarchical modeling of risk to sagebrush ecosystems from climate change, land use and invasive species in Nevada, USA. Ecography 33, 198208. https://doi.org/10.1111/j.1600-0587.2009.05684.x

Bradley, B.A., 2009. Regional analysis of the impacts of climate change on cheatgrass invasion shows potential risk and opportunity. Global Change Biology 15, 196–208. https://doi.org/10.1111/j.1365-2486.2008.01709.x

Bradley, B.A., Curtis, C.A., Fusco, E.J., Abatzoglou, J.T., Balch, J.K., Dadashi, S., Tuanmu, M.-N., 2018. Cheatgrass (Bromus tectorum) distribution in the intermountain Western United States and its relationship to fire frequency, seasonality, and ignitions. Biological Invasions 20, 1493–1506. https://doi.org/10.1007/s10530-017-1641-8

Brummer, T.J., Taylor, K.T., Rotella, J., Maxwell, B.D., Rew, L.J., Lavin, M., 2016. Drivers of Bromus tectorum Abundance in the Western North American Sagebrush Steppe. Ecosystems 19, 986–1000. https://doi.org/10.1007/s10021-016-9980-3

Buisson, L., Thuiller, W., Casajus, N., Lek, S., Grenouillet, G., 2010. Uncertainty in ensemble forecasting of species distribution. Global Change Biology 16, 1145–1157. https://doi.org/10.1111/j.1365-2486.2009.02000.x

Bureau of Land Management, 2016. About: Our Mission [WWW Document]. About: Our Mission. URL https://www.blm.gov/about/our-mission (accessed 4.29.19).

Camp, M.J., Rachlow, J.L., Shipley, L.A., Johnson, T.R., Bockting, K.D., 2014. Grazing in sagebrush rangelands in western North America: implications for habitat quality for a sagebrush specialist, the pygmy rabbit. The Rangeland Journal 36, 151. https://doi.org/10.1071/RJ13065

Chambers, J.C., Pyke, D.A., Maestas, J.D., Pellant, M., Boyd, C.S., Campbell, S.B., Espinosa, S., Havlina, D.W., Mayer, K.E., Wuenschel, A., 2014. Using resistance and resilience concepts to reduce impacts of invasive annual grasses and altered fire regimes on the sagebrush ecosystem and greater sage-grouse: A strategic multi-scale approach. Gen. Tech. Rep. RMRS-GTR-326. Fort Collins, CO: U.S. Department of Agriculture, Forest Service, Rocky Mountain Research Station. 73 p. 326. https://doi.org/10.2737/RMRS-GTR-326

Cole, K.L., Ironside, K.E., Arundel, S.T., Duffy, P., Shaw, J., 2008. Modeling future plant distributions on the Colorado Plateau: An example using *Pinus edulis*. In: van Riper, C. III; Sogge, M. K., eds. The Colorado Plateau III. Integrating Research and Resources Management for Effective Conservation. Tucson, AZ: The University of Arizona Press. p. 319–330.

Compagnoni, A., Adler, P.B., 2014. Warming, competition, and *Bromus tectorum* population growth across an elevation gradient. Ecosphere 5, art121. https://doi.org/10.1890/ES14-00047.1

Creutzburg, M.K., Halofsky, J.E., Halofsky, J.S., Christopher, T.A., 2015. Climate Change and Land Management in the Rangelands of Central Oregon. Environmental Management 55, 43–55. https://doi.org/10.1007/s00267-014-0362-3

Crimmins, S.M., Dobrowski, S.Z., Mynsberge, A.R., 2013. Evaluating ensemble forecasts of plant species distributions under climate change. Ecological Modelling 266, 126–130. https://doi.org/10.1016/j.ecolmodel.2013.07.006

Davies, G.M., Bakker, J.D., Dettweiler-Robinson, E., Dunwiddie, P.W., Hall, S.A., Downs, J., Evans, J., 2012. Trajectories of change in sagebrush steppe vegetation communities in relation to multiple wildfires. Ecological Applications 22, 1562–1577. https://doi.org/10.1890/10-2089.1

Deser, C., Phillips, A., Bourdette, V., Teng, H., 2012. Uncertainty in climate change projections: the role of internal variability. Clim Dyn 38, 527–546. https://doi.org/10.1007/s00382-010-0977-x

Dessai, S., Hulme, M., 2007. Assessing the robustness of adaptation decisions to climate change uncertainties: A case study on water resources management in the East of England. Global Environmental Change 17, 59–72. https://doi.org/10.1016/j.gloenvcha.2006.11.005

Dormann, C.F., Schymanski, S.J., Cabral, J., Chuine, I., Graham, C., Hartig, F., Kearney, M., Morin, X., Römermann, C., Schröder, B., Singer, A., 2012. Correlation and process in species distribution models: bridging a dichotomy. Journal of Biogeography 39, 2119–2131. https://doi.org/10.1111/j.1365-2699.2011.02659.x

Diniz-Filho, J.A.F., Mauricio Bini, L., Fernando Rangel, T., Loyola, R.D., Hof, C., Nogués-Bravo, D., Araújo, M.B., 2009. Partitioning and mapping uncertainties in ensembles of forecasts of species turnover under climate change. Ecography 32, 897–906. https://doi.org/10.1111/j.1600-0587.2009.06196.x

Elith, J., Leathwick, J.R., 2009. Species Distribution Models: Ecological Explanation and Prediction Across Space and Time. Annual Review of Ecology, Evolution, and Systematics 40, 677–697. https://doi.org/10.1146/annurev.ecolsys.110308.120159

Eyring, V., Bony, S., Meehl, G.A., Senior, C.A., Stevens, B., Stouffer, R.J., Taylor, K.E., 2016. Overview of the Coupled Model Intercomparison Project Phase 6 (CMIP6) experimental design and organization. Geoscientific Model Development 9, 1937–1958. https://doi.org/10.5194/gmd-9-1937-2016

Friend, A.D., Lucht, W., Rademacher, T.T., Keribin, R., Betts, R., Cadule, P., Ciais, P., Clark, D.B., Dankers, R., Falloon, P.D., Ito, A., Kahana, R., Kleidon, A., Lomas, M.R., Nishina, K., Ostberg, S., Pavlick, R., Peylin, P., Schaphoff, S., Vuichard, N., Warszawski, L., Wiltshire, A., Woodward, F.I., 2014. Carbon residence time dominates uncertainty in terrestrial vegetation responses to future climate and atmospheric CO 2. Proceedings of the National Academy of Sciences 111, 3280–3285. https://doi.org/10.1073/pnas.1222477110

Griffies, S.M., Danabasoglu, G., Durack, P.J., Adcroft, A.J., Balaji, V., Böning, C.W., Chassignet, E.P., Curchitser, E., Deshayes, J., Drange, H., Fox-Kemper, B., Gleckler, P.J., Gregory, J.M., Haak, H., Hallberg, R.W., Heimbach, P., Hewitt, H.T., Holland, D.M., Ilyina, T., Jungclaus, J.H., Komuro, Y., Krasting, J.P., Large, W.G., Marsland, S.J., Masina, S., McDougall, T.J., Nurser, A.J.G., Orr, J.C., Pirani, A., Qiao, F., Stouffer, R.J., Taylor, K.E., Treguier, A.M., Tsujino, H., Uotila, P., Valdivieso, M., Wang, Q., Winton, M., Yeager, S.G., 2016. OMIP contribution to CMIP6: experimental and diagnostic protocol for the physical component of the Ocean Model Intercomparison Project. Geoscientific Model Development 9, 3231–3296. https://doi.org/10.5194/gmd-9-3231-2016

Havstad, K.M., Peters, D.P.C., Skaggs, R., Brown, J., Bestelmeyer, B., Fredrickson, E., Herrick, J., Wright, J., 2007. Ecological services to and from rangelands of the United States. Ecological Economics 64, 261–268. https://doi.org/10.1016/j.ecolecon.2007.08.005

Hijmans, R.J., 2018. raster: Geographic Data Analysis and Modeling. R package version 2.8-4. https://CRAN.R-project.org/package=raster

Hufkens, K., Keenan, T.F., Flanagan, L.B., Scott, R.L., Bernacchi, C.J., Joo, E., Brunsell, N.A., Verfaillie, J., Richardson, A.D., 2016. Productivity of North American grasslands is increased under future climate scenarios despite rising aridity. Nature Climate Change 6, 710–714. https://doi.org/10.1038/nclimate2942

IPCC, 2007. Summary for policymakers, in: Parry, M.L., Canziani, O.F., Palutikof, J.P., van der Linden, P.J., Hanson, C.E. (Eds.), Climate Change 2007: Impacts, Adaptation and Vulnerability. Contribution of Working Group II to the Fourth Assessment Report of the Intergovernmental Panel on Climate Change. Cambridge University Press, Cambridge, UK, pp. 7–22.

Izaurralde, R.C., Thomson, A.M., Morgan, J.A., Fay, P.A., Polley, H.W., Hatfield, J.L., 2011. Climate Impacts on Agriculture: Implications for Forage and Rangeland Production. Agronomy Journal 103, 371. https://doi.org/10.2134/agroni2010.0304

Jiang, X., Rauscher, S.A., Ringler, T.D., Lawrence, D.M., Williams, A.P., Allen, C.D., Steiner, A.L., Cai, D.M., McDowell, N.G., 2013. Projected Future Changes in Vegetation in Western North America in the Twenty-First Century. Journal of Climate 26, 3671–3687. https://doi.org/10.1175/JCLI-D-12-00430.1

Johnsen, K., Samuelson, L., Teskey, R., McNulty, S., Fox, T., 2001. Process Models as Tools in Forestry Research and Management 7.

Kleinhesselink, A.R., Adler, P.B., 2018. The response of big sagebrush (*Artemisia tridentata*) to interannual climate variation changes across its range. Ecology 99, 1139–1149. https://doi.org/10.1002/ecy.2191

Knapp, P.A., 1996. Cheatgrass (Bromus tectorum L) dominance in the Great Basin Desert: History. persistence. and influences to human activities. Global Environmental Change 6, 37–52. https://doi.org/10.1016/0959-3780(95)00112-3

Knutti, R., 2008. Should we believe model predictions of future climate change? Philosophical Transactions of the Royal Society A: Mathematical, Physical and Engineering Sciences 366, 4647–4664. https://doi.org/10.1098/rsta.2008.0169

Krysanova, V., Hattermann, F.F., 2017. Intercomparison of climate change impacts in 12 large river basins: overview of methods and summary of results. Climatic Change 141, 363–379. https://doi.org/10.1007/s10584-017-1919-y

Larocque, G.R., Komarov, A., Chertov, O., Shanin, Liu, J., Bhatti, J.S., Wang, W., Peng, C., Shugart, H.H., Xi, W., Holm, J.A., 2015. Process-Based Model□: A Synthesis of Models and Applications to Address Environmental and Management Issues *.

Larson, C.D., Lehnhoff, E.A., Noffsinger, C., Rew, L.J., 2018. Competition between cheatgrass and bluebunch wheatgrass is altered by temperature, resource availability, and atmospheric CO2 concentration. Oecologia 186, 855–868. https://doi.org/10.1007/s00442-017-4046-6

Littell, J.S., McKenzie, D., Kerns, B.K., Cushman, S., Shaw, C.G., 2011. Managing uncertainty in climate-driven ecological models to inform adaptation to climate change. Ecosphere 2, art102. https://doi.org/10.1890/ES11-00114.1

Liu, Z., Wimberly, M.C., 2016. Direct and indirect effects of climate change on projected future fire regimes in the western United States. Science of The Total Environment 542, 65–75. https://doi.org/10.1016/j.scitotenv.2015.10.093

McDowell, N.G., Williams, A.P., Xu, C., Pockman, W.T., Dickman, L.T., Sevanto, S., Pangle, R., Limousin, J., Plaut, J., Mackay, D.S., Ogee, J., Domec, J.C., Allen, C.D., Fisher, R.A., Jiang, X., Muss, J.D., Breshears, D.D., Rauscher, S.A., Koven, C., 2016. Multi-scale predictions of massive conifer mortality due to chronic temperature rise. Nature Climate Change 6, 295–300. https://doi.org/10.1038/nclimate2873

Mouquet, N., Lagadeuc, Y., Devictor, V., Doyen, L., Duputié, A., Eveillard, D., Faure, D., Garnier, E., Gimenez, O., Huneman, P., Jabot, F., Jarne, P., Joly, D., Julliard, R., Kéfi, S., Kergoat, G.J., Lavorel, S., Gall, L.L., Meslin, L., Morand, S., Morin, X., Morlon, H., Pinay, G., Pradel, R., Schurr, F.M., Thuiller, W., Loreau, M., 2015. Predictive ecology in a changing world. Journal of Applied Ecology 52, 1293–1310. https://doi.org/10.1111/1365-2664.12482

Murphy, B.P., Yocom, L.L., Belmont, P., 2018. Beyond the 1984 Perspective: Narrow Focus on Modern Wildfire Trends Underestimates Future Risks to Water Security. Earth’s Future 6, 1492–1497. https://doi.org/10.1029/2018EF001006

Norberg, A., Abrego, N., Blanchet, F.G., Adler, F.R., Anderson, B.J., Anttila, J., Araújo, M.B., Dallas, T., Dunson, D., Elith, J., Foster, S.D., Fox, R., Franklin, J., Godsoe, W., Guisan, A., O’Hara, B., Hill, N.A., Holt, R.D., Hui, F.K.C., Husby, M., Kålås, J.A., Lehikoinen, A., Luoto, M., Mod, H.K., Newell, G., Renner, I., Roslin, T., Soininen, J., Thuiller, W., Vanhatalo, J., Warton, D., White, M., Zimmermann, N.E., Gravel, D., Ovaskainen, O., 2019. A comprehensive evaluation of predictive performance of 33 species distribution models at species and community levels. Ecological Monographs e01370. https://doi.org/10.1002/ecm.1370

Notaro, M., Mauss, A., Williams, J.W., 2012. Projected vegetation changes for the American Southwest: combined dynamic modeling and bioclimatic-envelope approach. Ecological Applications 22, 1365–1388. https://doi.org/10.1890/11-1269.1

Parmesan, C., 2006. Ecological and Evolutionary Responses to Recent Climate Change. Annual Review of Ecology, Evolution, and Systematics 37, 637–669. https://doi.org/10.1146/annurev.ecolsys.37.091305.110100

Parmesan, C., Yohe, G., 2003. A globally coherent fingerprint of climate change impacts across natural systems. Nature 421, 37–42. https://doi.org/10.1038/nature01286

Polley, H.W., Briske, D.D., Morgan, J.A., Wolter, K., Bailey, D.W., Brown, J.R., 2013. Climate Change and North American Rangelands: Trends, Projections, and Implications. Rangeland Ecology & Management 66, 493–511. https://doi.org/10.2111/REM-D-12-00068.1

Prudencio, L., Choi, R., Esplin, E., Ge, M., Gillard, N., Haight, J., Belmont, P., Flint, C., 2018. The Impacts of Wildfire Characteristics and Employment on the Adaptive Management Strategies in the Intermountain West. Fire 1, 46. https://doi.org/10.3390/fire1030046

Queirós, A.M., Huebert, K.B., Keyl, F., Fernandes, J.A., Stolte, W., Maar, M., Kay, S., Jones, M.C., Hamon, K.G., Hendriksen, G., Vermard, Y., Marchal, P., Teal, L.R., Somerfield, P.J., Austen, M.C., Barange, M., Sell, A.F., Allen, I., Peck, M.A., 2016. Solutions for ecosystem-level protection of ocean systems under climate change. Global Change Biology 22, 3927–3936. https://doi.org/10.1111/gcb.13423

R Core Team, 2018. R: A Language and Environment for Statistical Computing. R Foundation for Statistical Computing, Vienna, Austria.

Redmond, M.D., Cobb, N.S., Miller, M.E., Barger, N.N., 2013. Long-term effects of chaining treatments on vegetation structure in piñon–juniper woodlands of the Colorado Plateau. Forest Ecology and Management 305, 120–128. https://doi.org/10.1016/j.foreco.2013.05.020

Reeves, M.C., Bagne, K.E., Tanaka, J., 2017. Potential Climate Change Impacts on Four Biophysical Indicators of Cattle Production from Western US Rangelands. Rangeland Ecology & Management 70, 529–539. https://doi.org/10.1016/j.rama.2017.02.005

Rehfeldt, G.E., Crookston, N.L., Sáenz-Romero, C., Campbell, E.M., 2012. North American vegetation model for land-use planning in a changing climate: a solution to large classification problems. Ecological Applications 22, 119–141. https://doi.org/10.1890/11-0495.1

Reich, P.B., Hobbie, S.E., Lee, T.D., Pastore, M.A., 2018. Unexpected reversal of C _3_ versus C _4_ grass response to elevated CO _2_ during a 20-year field experiment. Science 360, 317–320. https://doi.org/10.1126/science.aas9313

Renwick, K.M., Curtis, C., Kleinhesselink, A.R., Schlaepfer, D., Bradley, B.A., Aldridge, C.L., Poulter, B., Adler, P.B., 2018. Multi-model comparison highlights consistency in predicted effect of warming on a semi-arid shrub. Global Change Biology 24, 424–438. https://doi.org/10.1111/gcb.13900

Robinson, D., Beukema, S., Greig, L., 2008. Vegetation models and climate change. https://doi.org/10.13140/2.1.2485.6327

Schlaepfer, D.R., Lauenroth, W.K., Bradford, J.B., 2012. Effects of ecohydrological variables on current and future ranges, local suitability patterns, and model accuracy in big sagebrush. Ecography 35, 374–384. https://doi.org/10.1111/j.1600-0587.2011.06928.x

Smith, 2017. Ternary: An R Package for Creating Ternary Plots. Zenodo, https://doi.org/10.5281/zenodo.2641809

Snyder, K.A., Evers, L., Chambers, J.C., Dunham, J., Bradford, J.B., Loik, M.E., 2019. Effects of Changing Climate on the Hydrological Cycle in Cold Desert Ecosystems of the Great Basin and Columbia Plateau. Rangeland Ecology & Management 72, 1–12. https://doi.org/10.1016/j.rama.2018.07.007

Still, S.M., Richardson, B.A., 2015. Projections of Contemporary and Future Climate Niche for Wyoming Big Sagebrush (Artemisia tridentata subsp. wyomingensis): A Guide for Restoration. Natural Areas Journal 35, 30–44.

Tittensor, D.P., Eddy, T.D., Lotze, H.K., Galbraith, E.D., Cheung, W., Barange, M., Blanchard, J.L., Bopp, L., Bryndum-Buchholz, A., Büchner, M., Bulman, C., Carozza, D.A., Christensen, V., Coll, M., Dunne, J.P., Fernandes, J.A., Fulton, E.A., Hobday, A.J., Huber, V., Jennings, S., Jones, M., Lehodey, P., Link, J.S., Mackinson, S., Maury, O., Niiranen, S., Oliveros-Ramos, R., Roy, T., Schewe, J., Shin, Y.-J., Silva, T., Stock, C.A., Steenbeek, J., Underwood, P.J., Volkholz, J., Watson, J.R., Walker, N.D., 2018. A protocol for the intercomparison of marine fishery and ecosystem models: Fish-MIP v1.0. Geoscientific Model Development 11, 1421–1442. https://doi.org/10.5194/gmd-11-1421-2018

Urban, M.C., 2015. Accelerating extinction risk from climate change. Science 348, 571–573. https://doi.org/10.1126/science.aaa4984

US EPA, O,. 2015. Level III and IV Ecoregions of the Continental United States [WWW Document]. US EPA. URL https://www.epa.gov/eco-research/level-iii-and-iv-ecoregions-continental-united-states (accessed 3.11.20).

U.S. Geological Survey, 2017. Federal Lands of the United States [WWW Document]. URL https://nationalmap.gov/small_scale/mld/fedlanp.html (accessed 9.12.19).

van Mantgem, P.J., Stephenson, N.L., Byrne, J.C., Daniels, L.D., Franklin, J.F., Fule, P.Z., Harmon, M.E., Larson, A.J., Smith, J.M., Taylor, A.H., Veblen, T.T., 2009. Widespread Increase of Tree Mortality Rates in the Western United States. Science 323, 521–524. https://doi.org/10.1126/science.1165000

Warszawski, L., Frieler, K., Huber, V., Piontek, F., Serdeczny, O., Schewe, J., 2014. The Inter-Sectoral Impact Model Intercomparison Project (ISI–MIP): Project framework. Proceedings of the National Academy of Sciences 111, 3228–3232. https://doi.org/10.1073/pnas.1312330110

Weisberg, P.J., Lingua, E., Pillai, R.B., 2007. Spatial Patterns of Pinyon–Juniper Woodland Expansion in Central Nevada. Rangeland Ecology & Management / Journal of Range Management Archives 60, 115–124.

Yapp, G., Walker, J., Thackway, R., 2010. Linking vegetation type and condition to ecosystem goods and services. Ecological Complexity 7, 292–301. https://doi.org/10.1016/j.ecocom.2010.04.008

Yates, K.L., Bouchet, P.J., Caley, M.J., Mengersen, K., Randin, C.F., Parnell, S., Fielding, A.H., Bamford, A.J., Ban, S., Barbosa, A.M., Dormann, C.F., Elith, J., Embling, C.B., Ervin, G.N., Fisher, R., Gould, S., Graf, R.F., Gregr, E.J., Halpin, P.N., Heikkinen, R.K., Heinänen, S., Jones, A.R., Krishnakumar, P.K., Lauria, V., Lozano-Montes, H., Mannocci, L., Mellin, C., Mesgaran, M.B., Moreno-Amat, E., Mormede, S., Novaczek, E., Oppel, S., Ortuño Crespo, G., Peterson, A.T., Rapacciuolo, G., Roberts, J.J., Ross, R.E., Scales, K.L., Schoeman, D., Snelgrove, P., Sundblad, G., Thuiller, W., Torres, L.G., Verbruggen, H., Wang, L., Wenger, S., Whittingham, M.J., Zharikov, Y., Zurell, D., Sequeira, A.M.M., 2018. Outstanding Challenges in the Transferability of Ecological Models. Trends in Ecology & Evolution 33, 790–802. https://doi.org/10.1016/j.tree.2018.08.001

Zelikova, T.J., Hufbauer, R.A., Reed, S.C., Wertin, T., Fettig, C., Belnap, J., 2013. Eco-evolutionary responses of *Bromus tectorum* to climate change: implications for biological invasions. Ecology and Evolution 3, 1374–1387. https://doi.org/10.1002/ece3.542

[dataset] Zimmer, S., Grosklos, G., Adler, P., Belmont, P. (2019). Agreement and uncertainty among climate change impact models: A synthesis of sagebrush steppe vegetation predictions, HydroShare, https://doi.org/10.4211/hs.3b420b738128411e8e1e11b38b83b5f1

Ziska, L.H., Reeves, J.B., Blank, B., 2005. The impact of recent increases in atmospheric CO2 on biomass production and vegetative retention of Cheatgrass (Bromus tectorum): implications for fire disturbance. Global Change Biology 11, 1325–1332. https://doi.org/10.1111/j.1365-2486.2005.00992.x

