## Supplemental Figures 1-2 for "Agreement and uncertainty among climate change impact models: A synthesis of sagebrush steppe vegetation projections"

### Supplementary

**Figure S1**: Example RGB legend, demonstrating where points with varying mean values are plotted. Point values are shown in the included table.

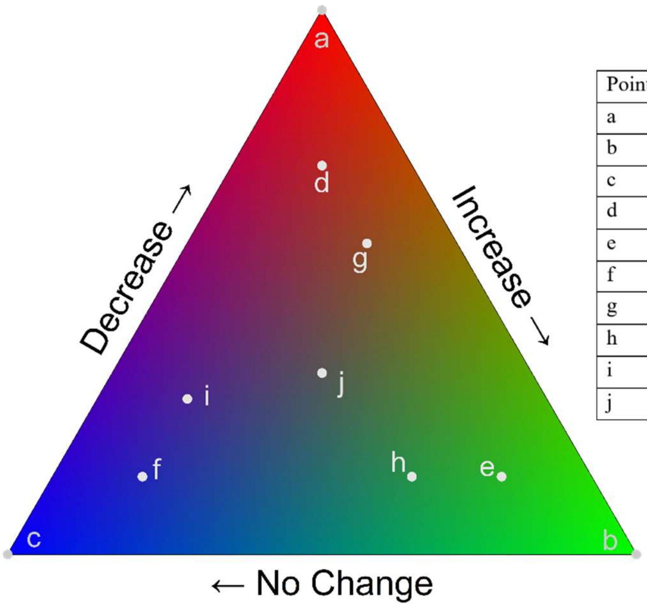


| Point | # Decrease | # Increase | # No change |
|---|---|---|---|
| a | 5 | 0 | 0 |
| b | 0 | 5 | 0 |
| c | 0 | 0 | 5 |
| d | 5 | 1 | 1 |
| e | 1 | 5 | 1 |
| f | 1 | 1 | 5 |
| g | 4 | 2 | 1 |
| h | 1 | 4 | 2 |
| i | 2 | 1 | 4 |
| j | 2 | 2 | 2 |

**Figure S2:** RGB visualizations for sagebrush results, showing only pixels that include non-gridded results from Renwick et al. 2018. Pixels are shown enlarged to increase visibility.

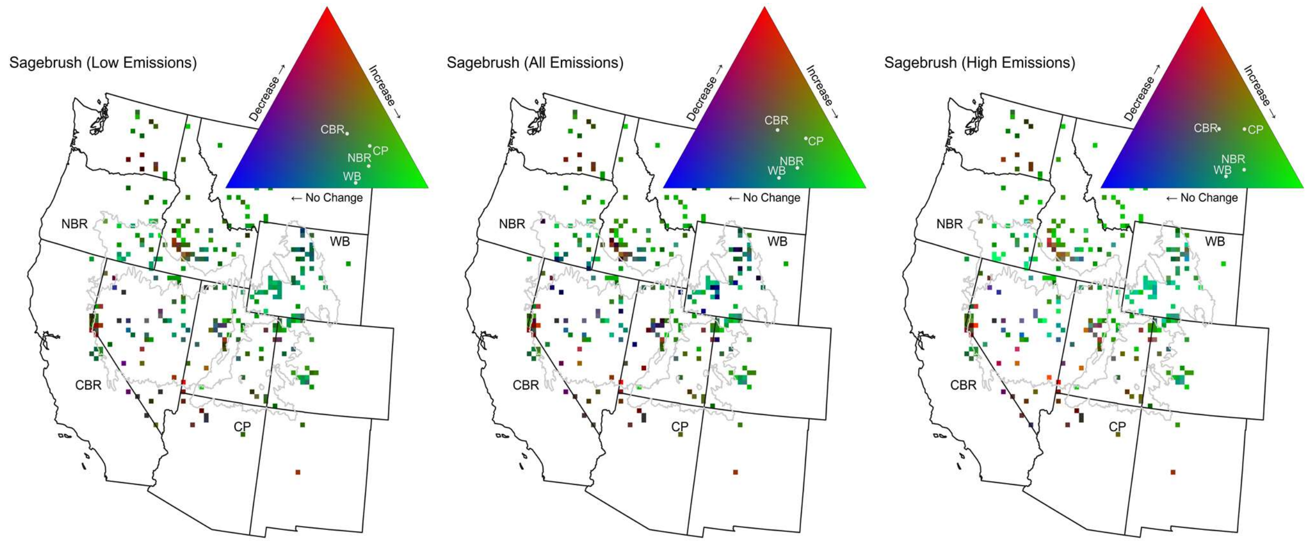


#### References

Renwick, K.M., Curtis, C., Kleinhesselink, A.R., Schlaepfer, D., Bradley, B.A., Aldridge, C.L., Poulter, B., Adler, P.B., 2018. Multi-model comparison highlights consistency in predicted effect of warming on a semi-arid shrub. Global Change Biology 24, 424–438. https://doi.org/10.1111/gcb.13900
